## Supplementary material for "Peptaibol Production and Characterization from *Trichoderma asperellum* and their Action as Biofungicide": Supplemeral table 1 and 2

Table S 1. Absolute and relative abundance of the most common amino acids in the sequence of P*aib* produced by *Trichoderma* species

| Aminoacid | Absolute abundance | Relative abundance (%) |
| --- | --- | --- |
| Aib | 1156 | 36.9 |
| Leucine/Isoleucine | 465 | 14.8 |
| Glutamine | 319 | 10.2 |
| Valine/Isovaline | 305 | 9.7 |
| Alanine | 282 | 9.0 |
| Proline | 261 | 8.3 |
| Glycine | 145 | 4.6 |
| Phenylalanine | 72 | 2.3 |
| Serine | 43 | 1.4 |
| Asparagine | 38 | 1.2 |
| Glutamate | 32 | 1.0 |
| Tryptophan | 10 | 0.3 |
| Others | 8 | 0.3 |
| Total | 3136 | 100 |

Source: based in the information reported at the P_aib_ Database [55]

Table S 2. Regression coefficients and probabilities associated with the factors in the model for predicting the production of P_aib_.

| Factor | Coeficient | Probability |
| --- | --- | --- |
| Constant | 4.05E+08 | <0.001* |
| Aib (A) | 4.28E+08 | 0.0002* |
| *F. oxysporum* (F) | -4.51E+08 | 0.7637 |
| Interaction (A-F) | -2.85E+07 | 0.4842 |
| Aib quadratic (A-A) | -5.78E+07 | 0.0887 |
| *F. oxysporum* quadratic (F-F) | 1.45E+08 | 0.0016* |

*Indicates the variable has a significative effect over the observed response (P < 0.05).
